# The function of environmentally acquired perfume blends in male orchid bees

**DOI:** 10.1101/2022.12.08.519619

**Authors:** Jonas Henske, Nicholas W. Saleh, Thomas Chouvenc, Santiago R. Ramírez, Thomas Eltz

## Abstract

Perfume making by male orchid bees is a remarkable behavior giving rise to an entire pollination syndrome in the neotropics. Male orchid bees concoct and store perfume mixtures in specialized hind-leg pockets using volatiles acquired from multiple environmental sources, including orchids. However, the precise function and ultimate causes of this behavior have remained elusive. We supplemented male *Euglossa dilemma* reared from trap-nests with perfume loads harvested from wild conspecifics. In dual-choice experiments, males supplemented with perfumes mated with more females, and sired more offspring, than untreated, equal-aged, control males. Our results demonstrate that male-acquired perfumes function as chemical signals emitted during courtship and received by females when selecting mates. Sexual selection might be a key agent shaping the evolution of perfume signaling.

**One-Sentence Summary:** The possession of exogenous volatiles increase male mating success and paternity in orchid bees.

## Main Text

The neotropical orchid bees are best known for the pollination services they provide to a large number of tropical plants, including hundreds of orchid species, thanks to the peculiar behavior in which male bees collect odoriferous substances from a broad range of sources in their environment (*1–3*). Male orchid bees store and accumulate volatile compounds in enlarged hind-leg pouches that facilitate the concoction of complex perfume blends (*4–6*). Ever since its original description by Darwin and others (*7–9*), this behavior has intrigued scientists and naturalists alike, but a major question remains unanswered: what is the purpose of the acquired substances?

The recipient of the chemical signal has remained unknown, but perfumes are believed to transfer chemical information emitted during courtship display. After accumulating perfume compounds, male bees actively release them during a stereotypical display performed at vertical stems (perches) in the forest understory (*4*) (Fig. 1A). Additionally, comparative studies have shown that perfumes have evolved exceptionally fast (*10*) into species-specific chemical blends that may facilitate mate recognition (*11*). Males display on the downwind side of the perch (*12*), and females have been observed to approach the male upwind and land at or near the perch, where mating takes place (*12–17*). Therefore, perfumes have been hypothesized to function as *inter*-sexual sex pheromone analogues that mediate mate recognition and/or indicate the genetic quality of males (*18–20*).

**Fig. 1.**
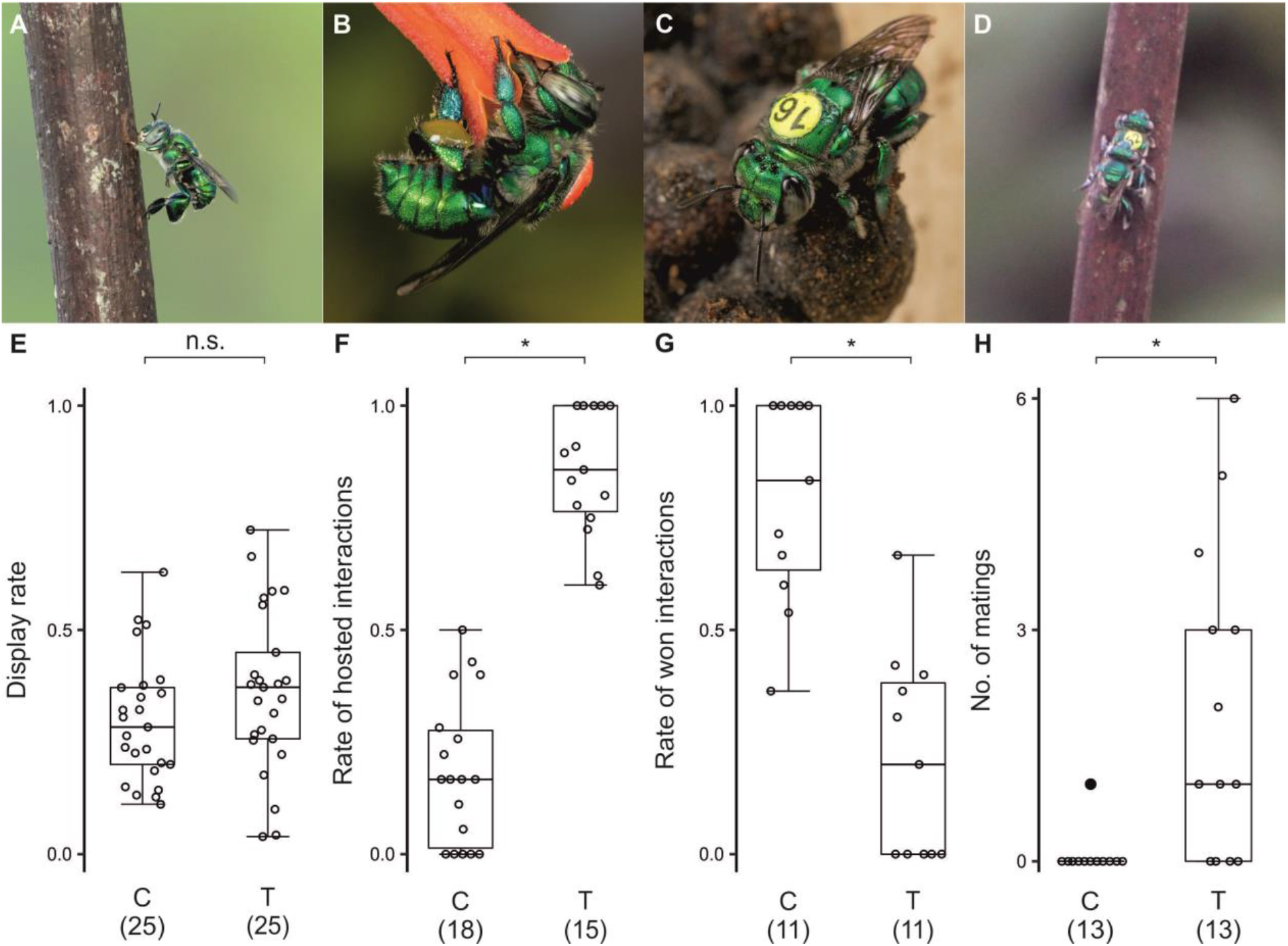
Reproductive success and perfume-dependent behaviors of *Euglossa dilemma* during flight cage experiments. **(A)** Male showing characteristic perching behavior during display. **(B)** Marked female with pollen loads drinking nectar from *Hamelia patens* flower. **(C)** Marked female on brood cells constructed in wooden nest box. **(D)** Copulation event during experiment after female (with tag) landed on the perch (image captured from video sequence, see movie S1). **(E)** Male display intensity did not vary with perfume supplementation (T). **(F)** Perfume-supplemented males received more visits (i.e. hosted more interactions) at their perch site by control males, whereas **(G)** control males (C) remained more often at contested perch following those interactions. **(H)** Successful mating events were achieved almost exclusively by perfume-supplemented males. **(E-H)** Numbers in brackets indicate sample size, C=control bees with no perfume, T=treatment bees supplemented with perfume. Asterisks (*) indicate significant differences at P<0.01, Mann-Whitney U. Boxplots show median (center line), upper and lower quartile (box limits), 1.5x interquartile range (whiskers), individual data points (unfilled dots) and outliers (black dots).

Experimental evidence of female attraction to perfumes is lacking, and bioassays with perfume extracts only attracted other conspecific males (*17*). When a male bee displays and holds a display territory, other conspecific males are often attracted to the area and—after a brief interaction— only one competitor retains the perch. These observations raise the possibility that perfumes may also transmit *intra*-sexual information about the competitive prowess of a male, thus resolving conflict either though usurpation or defense of courtship territories (*13, 16*). Equivalent visual signals have evolved across animals whereby both inter-sexual and intra-sexual communication is transmitted (*21, 22*). For example, the “skraa calls” of bower birds first evolved for aggressive display during male contests and were later co-opted for courtship and female choice (*23*). Similarly, orchid bee perfumes may play a role in both inter-sexual and intra-sexual communication, mediating mate choice and establishing dominance hierarchies among males.

To elucidate the function of male perfumes in mate choice, we conducted cage experiments with the orchid bee *Euglossa dilemma*, a Mesoamerican species that was recently naturalized in South Florida (*24, 25*). We obtained freshly hatched bees using trap-nests and we individually marked males and females and released them into two large flight cages equipped with perch sites for male display and floral resources for food and nesting materials. Females used resin to construct brood cells inside wooden boxes that they provisioned with pollen for larval consumption (Fig. 1B, C); males and females fed on nectar flowers. No more than two displaying males were present per cage at any time. We supplemented one male with 1μL of perfume harvested from wild conspecifics before introduction to the cage while the other male was handled in the same way but received no perfume supplement. In dual choice experiments, we tested the effect of perfume supplementation on male display activity, on the initiation and outcome of male-male interactions, and on male mating success (see supplementary methods).

Perfume supplementation and marking of male bees took place immediately after hatching, and both treatment and control bees were introduced to the cages the following day. To validate the effect of perfume supplementation, and to assess perfume dynamics during the experiment, we analyzed hind-leg extracts via gas chromatography and mass spectrometry (GC/MS). We collected and analyzed leg extracts from *i*) freshly hatched males, *ii*) males that had been supplemented with perfume on the day before, and *iii*) experimental males of both treatment and control groups upon sampling after they spent 10±3.6 (m±sd) days in the cage. Hind-leg pouches of freshly hatched bees did not contain any volatiles (N=9), whereas males supplemented with perfume the day before (N=8) had maximum perfume loads containing higher amounts (P<0.001) and more compounds (P=0.001) than residual perfumes of experimental males (N=24; Fig. S1). Males of the control group (N=20), however, were not completely empty, but managed to acquire some volatiles in the cage, possibly from leaves of food plants (J.H. pers. obs.) or from microbes associated with living or decaying plant parts(*3*). However, the overall quantity of their volatiles was substantially lower than that of treatment males (P<0.001, Fig. S1A). In addition, the volatiles found in control males were less complex (P<0.001, Fig. S1B) and different in chemical composition compared to those of treatment males (Fig. S2, S3). Thus, volatile compounds that were available in the cages did not reflect the diversity found in wild caught individuals. These analytical results show that perfume supplementation was effective in raising quantity and complexity of hind-leg volatiles in treatment males for the entire duration of each trial, consistent with (1) hind-leg pouches being efficient volatile storage containers (*26*) and (2) moderate loss of perfume contents during display by experimental males (*4*).

Males started to exhibit display behavior on average 3.1±2.1 days after being released into the cage with some individuals displaying on the first day. There was no difference neither in the onset of display (N=50; P=0.422) nor in display activity between experimental groups (N=50; P=0.123; Fig. 1E), suggesting that display behavior is highly stereotypic and does not require previous collection, or possession, of volatiles.

The possession of perfume loads had a strong influence on the initiation and the outcome of male-male interactions that regularly took place during display bouts. To the observer, these interactions appear competitive in nature, with males engaging in ritualized zig-zag or sustained circling flights near the perch. These interactions rarely involve physical contact and usually end with one of the males leaving the site and the other resuming display at the perch site where the encounter took place (*13–17*) (pers. obs.). Male *E. dilemma* regularly interacted in this manner during display in the experimental cages. Interactions took place more often at the perch of the treatment male (N=33, P<0.001; Fig. 1F). However, the possession of perfume did not increase the likelihood that a male retains his perch. To the contrary, the control male more frequently took over and resumed display at the interaction perch (N=22, P<0.001; Fig. 1G). These results refute the hypothesis that perfumes function as signals of competitive prowess in male-male competition for perch sites but suggest that control males were attracted by the perfume stimuli released by experimental males. This is congruent with field observations where male orchid bees (1) are attracted to conspecific perfumes in bioassays and (2) usually approach conspecific display sites from downwind (*17*). Our observations indirectly support the idea that males visit the display sites of other males as an opportunistic strategy for sneaking copulations (*13*), a low-cost strategy for males that have not yet acquired sufficient perfume to attract mates on their own (*13*). Our results are ambiguous with regard to the hypothesis of male orchid bees forming leks (*16*). Leks are non-resource based cooperative aggregations formed by males engaging in joint courtship to entice females (*27*). In the natural habitat orchid bees display sites are sometimes clumped in space, e.g. around treefall clearings or on top of hills and ridges, possibly representing an “exploded” lek (*12, 16*) That supplemented males were more frequently visited by the partner male is in general agreement with the idea of leks. Thus, perfumes could serve as a social signal among males to congregate, increasing the individual chance for copulation. Males might evaluate perfumes of conspecifics during interaction flights. Individuals without species-specific perfumes would not contribute in strengthening the social chemical signal. Therefore, treatment individuals might opt to escape from territorial interactions to claim a new territory or to seek other conspecifics that possess perfume. This would explain why treatment males leave the perch following the interactions more often than control males in our experiment.

Female orchid bees mate only once in their lifetime resulting in functional monogamy (*28*). Thus, male access to females is very limited and copulations are difficult to observe (*28*) even in a flight cage (but see below). To test if male mating success depends on the possession of species-specific perfume, we genotyped diploid female offspring using microsatellite markers. Each female mother and her corresponding larvae offspring were sampled after females had completed at least four brood cells. Dissections of spermathecae revealed sperm in 51.9 % of females, showing that they had mated in the cages. Paternity analysis unambiguously confirmed single mating in those females (*28*) and revealed that the brood of 26 out of 27 inseminated females was sired by the treatment males (P<0.001). Treatment males mated more often than control males (N=26; P=0.005) and had a higher individual mating success than control males (N=26; P=0.003; Fig. 1H). These results constitute the first direct evidence that the possession of male perfume mixtures modify the mating preference by female orchid bees. In fact, only one female mated with a control male in our experiment. Notably, this control male was exceptional in possessing a perfume similar in composition to the corresponding treatment male (Fig. S4). It is known that male orchid bees occasionally collect perfume from the hind tibial surface of other conspecific males, dead or alive, especially in captive conditions (*29*) (pers. obs.). Although we cannot rule out the possibility that the control male obtained the perfume before mating, it is likely that perfume theft facilitated this mating event.

In addition to inferring paternity via genotyping, we directly observed eight copulation events during the course of the experiment; in all cases, the female approached the male perch from downwind in a zigzag-flight pattern in accordance with anemotactic tracking of a perfume plume (*30*). In most cases, females quickly landed on the display perch underneath the male and mating took place, lasting only a few seconds (see Movie S1). In one case, the female flew back and forth between the two perch sites, closely inspecting the control male, but finally making a choice to copulate with the perfume male. These observations support the hypothesis that perfumes transfer information during pre-mating behavior.

## Conclusion

In this study, we demonstrate that female orchid bees select conspecific mates based on the possession of complex perfume mixtures. Our results showed that perfume directly affects male mating success through female choice, therefore supporting the hypothesis that perfumes are *inter*-sexual signals that transfer information during male courtship display. In addition, we showed that perfumes released by displaying males attract other males and facilitate the location of conspecific courtship territories.

Sex pheromones have been shown to mediate species recognition in a diverse array of insect lineages, including moths and beetles (*31–33*). Due to the strong importance on reproductive success, natural selection is thought to have optimized the recognition functions of sex pheromones (*18, 20*). In fact, previous comparative studies of perfume phenotypic diversity support the view that natural selection has shaped chemical specificity and divergence across lineages of orchid bees (*10, 11, 34*). However, natural selection is not the only force at play. For instance, recent studies revealed substantial heritable variation in pheromone traits, especially in male-calling systems, suggesting that sex pheromones can also evolve by sexual selection (*19, 35–38*). With perfume volatiles being scarce and unpredictable in natural habitats (*1, 39, 40*), the process of concocting perfume is likely costly to males, which provides an opportunity for perfumes to evolve as honest indicators of survival/age, foraging success, cognitive skill and/or competitive strength (*34, 39*). Under this scenario, females that respond correctly to perfume stimuli before mating are effectively selecting males that exhibit such fitness components. What exactly constitutes an attractive perfume needs to be addressed in future choice experiments. By manipulating parameters like perfume chemical composition and complexity, it may be possible to see how the presence of certain specific major compounds, like HNDB in *Euglossa dilemma* (*41*), contributes to perfume attractiveness. Mutations in genes of the olfactory system are known to affect HNDB perception in males (*42*), but they likely also affect perception in females, possibly resulting in concerted evolution of male signals and female preferences (*41*). This and previous studies support the idea that both natural selection and sexual selection shape perfume signaling in orchid bees.

## Acknowledgments

We thank the Fern Forest Nature Center, the Flamingo Gardens Wildlife Sanctuary and Joe Patt for the opportunity to place trap-nests. We also thank Aaron Mullins, Jorge Gomez and all the staff from the UF Fort Lauderdale Research and Education Center for their support. We are grateful to Philipp Brand for help with mining microsatellite loci, Maximilian Schweinsberg for assistance with the paternity analysis and to Tamara Pokorny for ideas and expertise regarding experimental design.

## Author contributions

Conceptualization: TE, SRR; Methodology: JH, NWS, SRR, TE; Investigation: JH, NWS; Formal analysis: JH; Visualization: JH; Resources: TC, SRR, TE; Funding acquisition: JH, SRR, TE, NWS; Supervision: TE; Writing – original draft: JH; Writing – review & editing: JH, TE, SRR, TC, NWS

## Data availability

Raw data supporting the findings of this study were uploaded to figshare and will be published before publication. Any data is available from the corresponding author upon request.

## List of Supplementary materials

Materials and Methods

Figs. S1 to S4

Table S1

References (*43–50*)

Images S1 to S3

Movies S1 to S3

