## Supplementary Material for "The function of environmentally acquired perfume blends in male orchid bees"

**This PDF file includes:**

Materials and Methods

Figs. S1 to S4

Table S1

Images S1 to S3

Captions for Movies S1 to S3

**Other Supplementary Materials for this manuscript include the following:**

Movies S1 to S3

Materials and Methods

Experiments took place in 2018, 2019 and 2021 (August – December) on the Campus of the University of Florida (UF) Ft. Lauderdale Research and Education Center in Davie, Florida. We used one cage in 2018 and 2019 (15 x 15 x 4 m, see Image S1) and two cages in 2021 (15 x 15 x 4 m; 9 x 9 x 3 m). Experimental flight cages were made from shade cloth to provide forest understory conditions. We obtained plants from local nurseries, mostly: *Hamelia patens* (nectar), *Penta sp*. (nectar) and *Senna alata* (pollen). We used stingless bee cerumen obtained from stingless bee colonies (*Melipona panamica*) in Panama as a main resin source. Wooden nest boxes were placed in the cages to facilitate nest building of females. We obtained experimental bees by trap-nesting bees in South Florida in wooden boxes placed around buildings on the campus of the UF Fort Lauderdale Research and Education Center, at Fern Forest Nature Center, Flamingo Gardens and at private residences. We checked nests daily for newly hatched bees to use for experimentation.

**Experimental setup**

We marked hatched females with numbered tags (Opalith-tags; Holtermann Imkereibedarf, Brockel, Germany) and introduced them to the experimental cages on their first day of adult life. We checked nest boxes in the cages every night and sampled females and their corresponding offspring after they had built and provisioned four brood cells. We immediately dissected spermathecae to examine mating status (*43*). Females and their brood cells were transferred to 95%-alcohol for later DNA extraction. We marked freshly hatched males with scratch marks on the thorax (*44*). Treatment bees received perfume supplementation (see below). Control bees were handled in the same way but did not receive perfume. Both control and treatment bees were kept individually for one night in small insectaria (40 x 40 x 20 cm) equipped with a nectar plant before releasing them into the experimental cages on the following day. The timing of introduction of experimental males in the cages varied somewhat depending on the availability of freshly hatched individuals, which sometimes was a limiting factor. Males were consecutively introduced ensuring the presence of a maximum of one control and one treatment male displaying in one cage at any given time. After sampling, males were conserved in 95%-alcohol. Hind-legs were sampled separately (see GC/MS analysis). Since pollen is a limited resource for breeding females, we did not allow more than 20 females to be active in a cage at any given time. In total 55 of 102 females were inseminated, of which we considered 27 for downstream paternity analysis. The remaining inseminated females had to be discarded because either the entire offspring consisted of haploid individuals or some experimental males were lost to unknown causes in the cage and thus could not be genotyped to ensure reliable paternity assignment between control and treatment males. Therefore, we discarded all data from mated females that could have been inseminated by the lost males. We emphasize that, in doing so, we did not discard any known copulations of control males. In total 26 males (13 control and 13 treatment males) were included in the paternity analysis. These males spent an average of 10.7±3 (m±sd) days in the cage with no difference between experimental groups (P=0.687). For analysis of perfume content in male hind legs and behavioral differences between experimental groups (see below), all sampled or observed males were taken into account (including some that were lost and excluded from paternity analysis).

**Behavioral observations**

We monitored male display and interaction behavior using standardized 10-min observation intervals during which we scanned all potential perch sites, i.e. vertical stems and structures, three times. Monitoring took place during 5 - 10 observation intervals per day between 7am and 11:30am, when bees were most active. We identified displaying males using binoculars (Pentax Papilio) to recognize specific scratch marks on the thoraces of the treatment and control bees. For each male, we quantified overall display activity (likelihood to display per interval averaged over all 10-min intervals), the frequency of hosting male interactions (received interactive visits from the other male) at display perches, and the outcome of such interactions. An outcome of an interaction was only recorded when one male retreated from the interaction and the other one remained at the perch site resuming display. For interaction analyses, we only considered males, which had participated in at least five interactions. Males that never showed display (N=4) were excluded from the analysis.

**Perfume supplementation**

To supplement experimental males with species-specific perfume we attracted wild *Euglossa dilemma* at Fern Forest Nature Center using screened p-dimethoxybenzene baits. Bees were caught with a hand net and we harvested perfume by gently squeezing the hind-legs of the bees absorbing the perfume with 2µL-microcapillary tubes (Hirschmann Laborgeräte GmbH, Eberstadt, Germany; see Movie S2). The extracted perfumes were stored in glass-vials at -20°C. Each perfume we used to supplement experimental males consisted of extracted perfumes of 5 to 8 wild-caught individuals, depending on yield. For perfume supplementation 0.5µL of the harvested perfume were applied to each hind-leg pocket of treatment males (for further technical information see (*45*)). To apply harvested perfumes we used 2µL-microcapillary tubes in 2018 and 2019 and a 1.2µL-CTC-syringe (Hamilton, Inc, Reno, NV, USA) in 2021 (see Movie S3).

**Paternity analysis**

New microsatellite markers were designed with Geneious Prime (Dotmatics, Boston, MA, USA) software using the genome of *Euglossa dilemma* (*(46);*see Table S1). We extracted DNA from sampled males, females and brood using DNeasy Blood and Tissue kit (QIAGEN, Inc, Valencia, CA, USA) following the manufacturer’s protocol. Multiplex-PCR was conducted using GoTaq (Promega, Inc, Madison, WI, USA) and consisted of 30 cycles of 94° for 30s, 60°for 90s and 72° for 90s with an initial step of 95° for 2min and a final elongation step of 72° for 10min. Forward primers were labeled with fluorescent tags (6-FAM, ATTO532, ATTO550, ATTO565). Fluorescent PCR products were then diluted with water to a 1:10 ratio and combined with HiDi formamide and Liz 500 size (in 2021 Liz 600) standard. Fragment sizes were measured with an ABI 3730XL DNA analyzer (Applied Biosystems, Inc, Foster City, CA, USA) at Microsynth SeqLab in Switzerland. Alleles were identified using GeneMarker (Softgenetics LLC, State College, State College, PA, USA) software and we determined paternity by visually comparing alleles among parents and offspring taking advantage of the haplodiploid reproduction system of orchid bees.

**GC/MS analysis**

We analyzed perfume volatiles in male hind-legs from the following males: freshly hatched males, males that had been supplemented with perfume on the day before, and experimental males of both treatment and control groups at the end of their cage time. We transferred hind-legs to vials containing 500µL of n-hexane for GC/MS analysis. Samples were stored at -20°C until chemical analysis in Bochum, Germany. Samples were analyzed using a HP5890II gas chromatograph coupled to a HP5972 mass spectrometer (Hewlett-Packard, Palo Alto, CA, USA), equipped with a DB- 5MS column (30 m, 0.25 μm film thickness, 0.25 mm diameter), with splitless injection (2μL). The GC oven was programmed from 60 to 300°C at 10°C/min followed by 15 min isothermal at 300°C. For further analysis cuticular hydrocarbons and long chain alcohols and acetates known to derive from bees’ labial glands were excluded (*5, 47, 48*). We analyzed perfume complexity (number of compounds) and perfume quantity (summed peak area of all compounds) using the software ChemStation (Agilent Technologies, Santa Clara, CA, USA). We identified compounds using commercial mass spectral libraries (*49, 50*) in conjunction with our own user libraries. We calculated the relative abundance of each compound relative to the total amount of perfume for control and treatment bees.

**Statistics**

We tested for differences in display intensity, onset of display behavior, time spent in the cages, the frequency of hosting male interactions, the outcome of male interactions and individual mating success between control and treatment males using two-tailed Mann-Whitney U tests as implemented in SPSS statistics v. 28.0.0.0 (IBM, Armonk, NY, USA). We used the same software to test for differences in perfume quantity and complexity between control and treatment and treatment and freshly hatched perfume-supplemented males using two-tailed Mann-Whitney U tests. We used Yates modified two-tailed chi-square test to test for difference in the proportion of treatment and control males contributing to copulations. To test whether the total number of successful copulations depended on treatment we used a two-tailed binominal test. The data were recorded with Microsoft Excel (v.16.48). The figures were plotted in R (v.4.2.1).

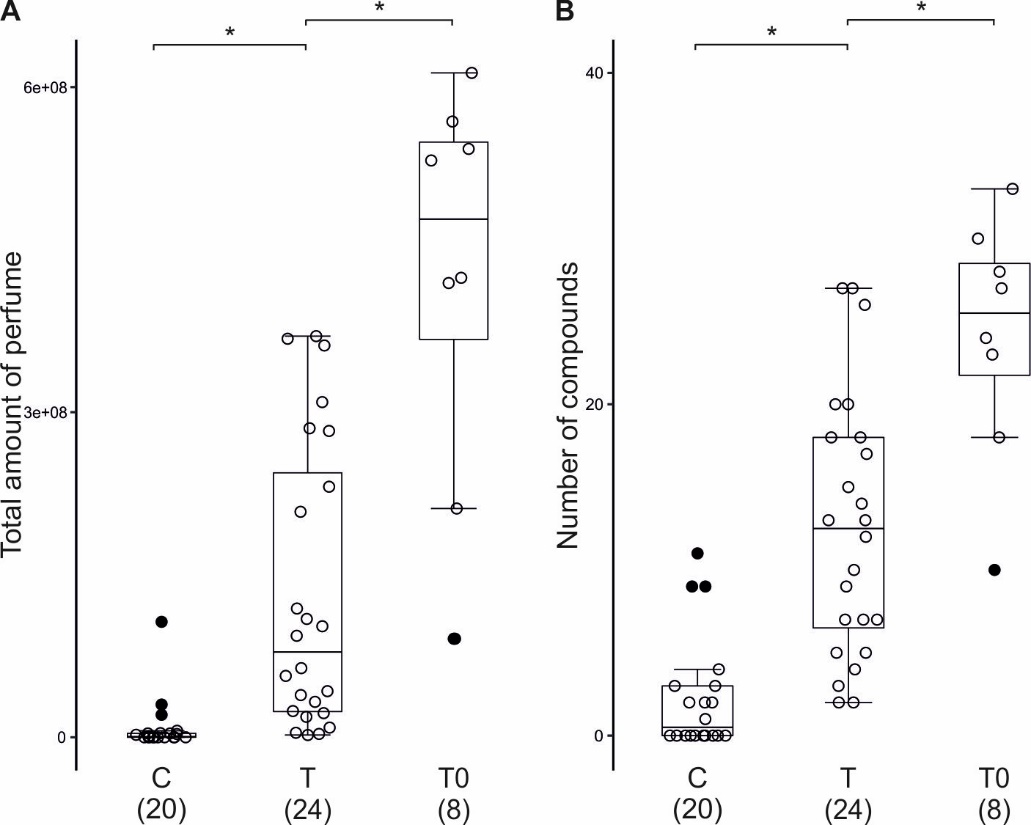

Fig. S1.

**Comparisons of perfume contents between experimental groups** **(A)** Total amount of perfume found in hind-legs. **(B)** Number of compounds of analyzed perfumes. **A,B** Numbers in brackets indicate sample size, C=control bees without perfume supplementation, T=Experimental bees supplemented with species-specific perfume and sampled at the end of the trial, T0=Perfume-supplemented bees sampled after one day. Freshly hatched bees without perfume-supplement (N=9) sampled after one day did not contain any perfume and are not shown. Asterisks (*) indicate significant differences at P<0.01, Mann-Whitney U. Boxplots show median (center line), upper and lower quartile (box limits), 1.5x interquartile range (whiskers), individual data points (unfilled dots) and outliers (black dots).

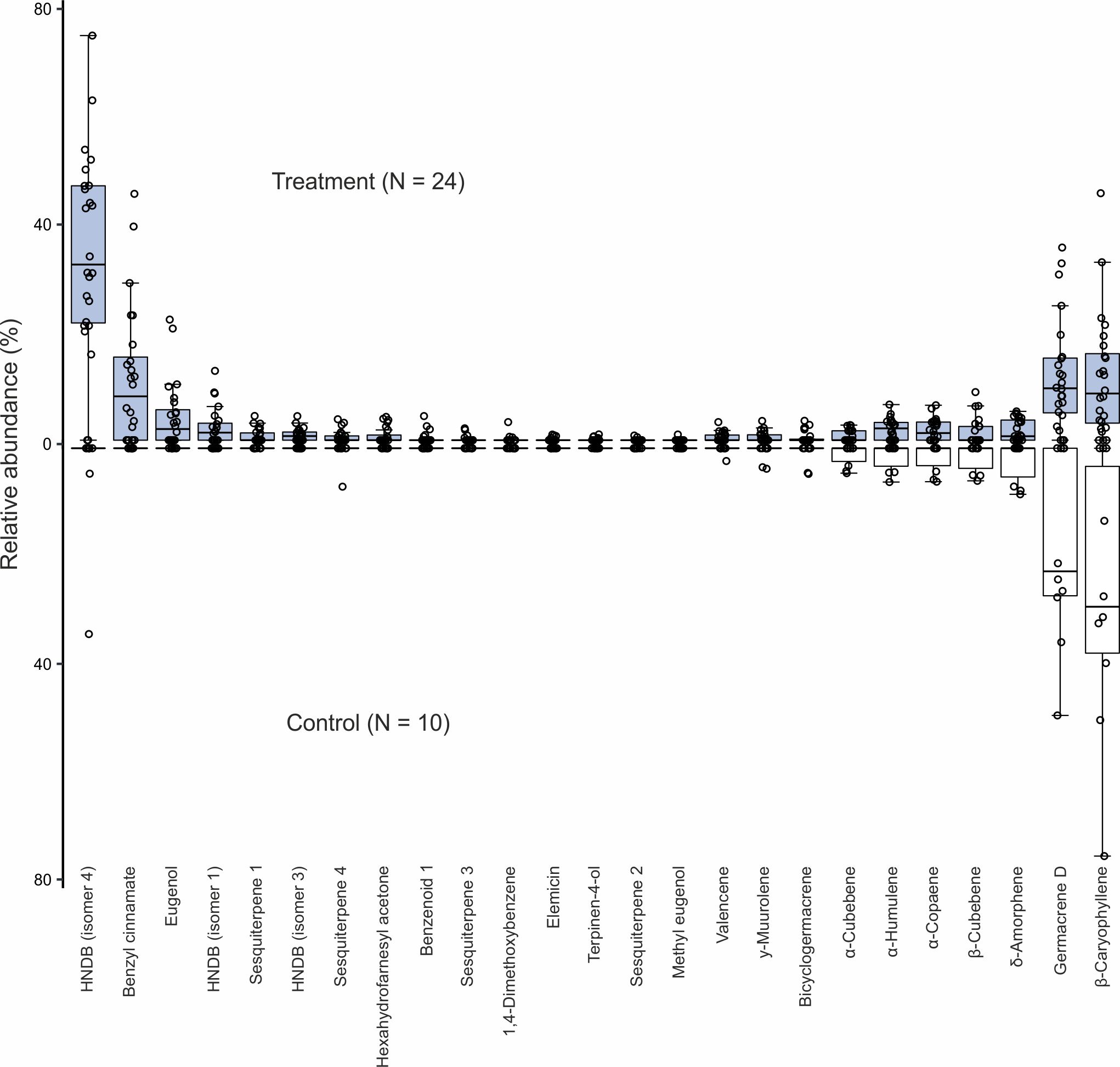

Fig. S2.

**Composition of perfumes of experimental males (relative abundances)**. The 25 most abundant compounds are shown. Boxplots represent relative abundances (average percentage contribution of total peak area). Boxplots show median (center line), upper and lower quartile (box limits), 1.5x interquartile range (whiskers) and individual data points (dots). Experimental control males which contained no perfume (N=10) are not shown.

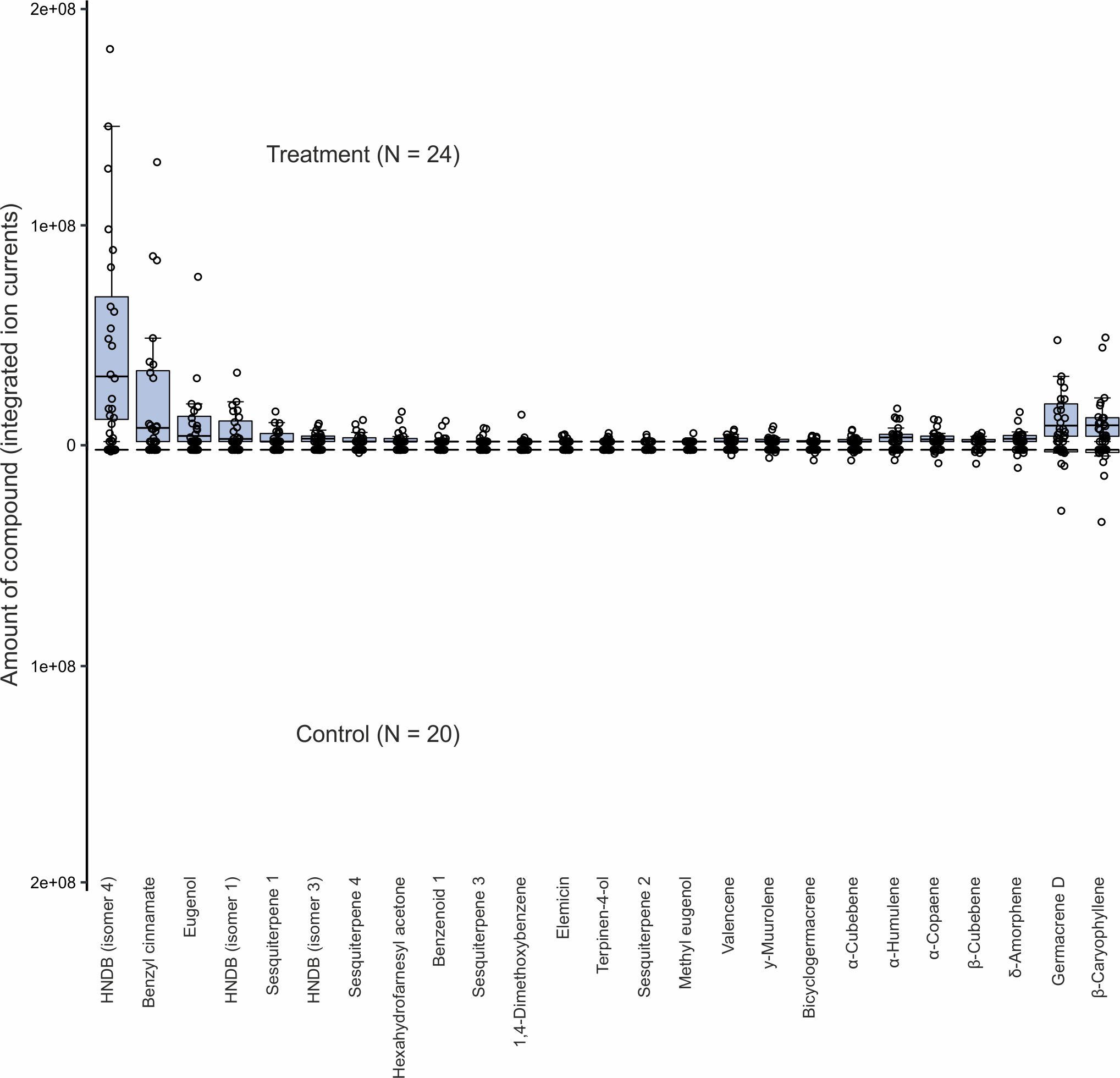

Fig. S3.

Composition of perfumes of experimental males (total amounts). The total amounts (integrated ion currents) of the 25 most abundant compounds are shown. Boxplots show median (center line), upper and lower quartile (box limits), 1.5x interquartile range (whiskers) and individual data points (dots).

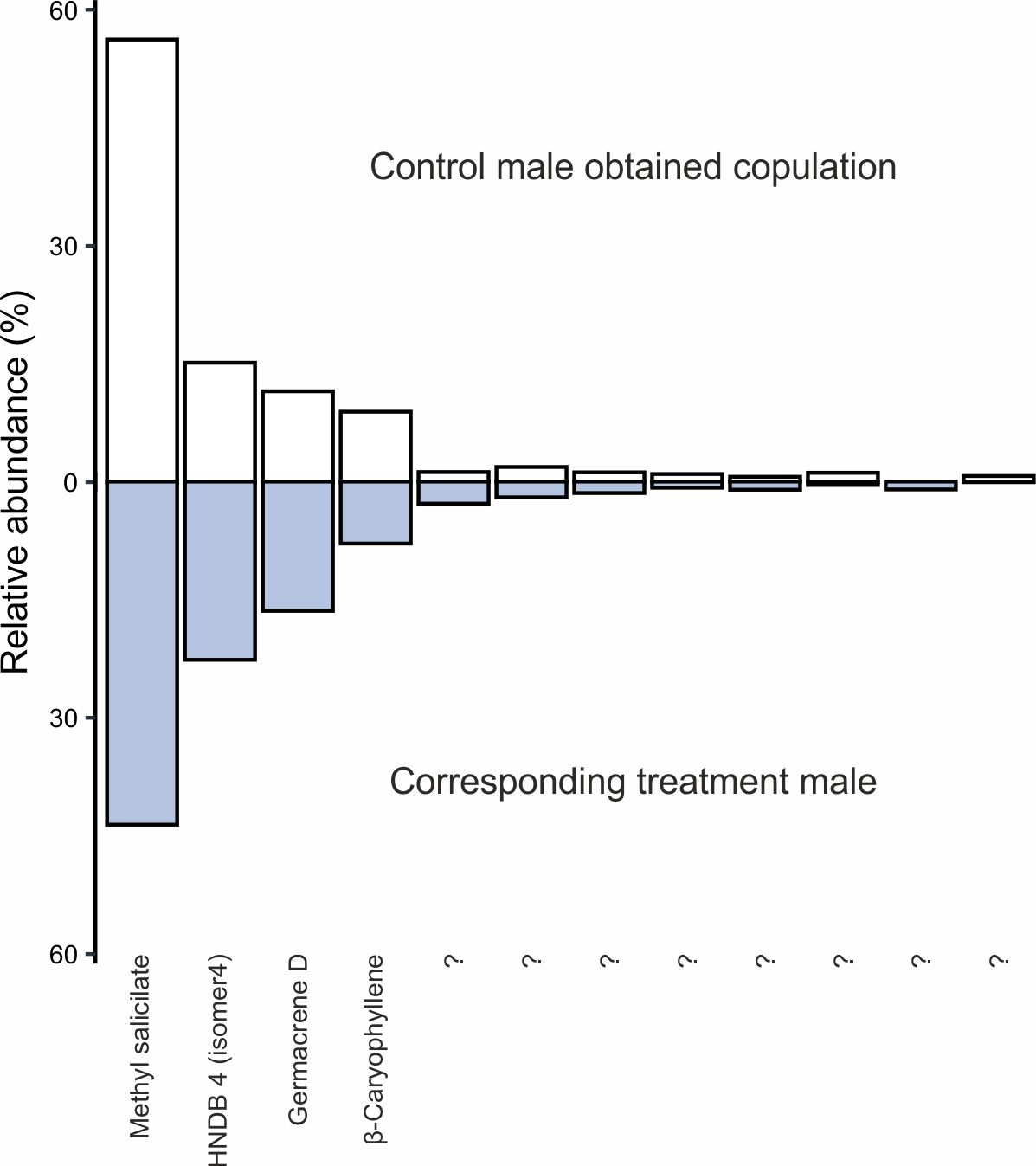

Fig. S4.

**Composition of perfumes of the control male that obtained one copulation and its corresponding treatment male**. All the compounds present in perfumes are shown. Bars represent relative abundances (percentage contribution of total peak area). Compound names are given when available.

Table S1.

*E. dilemma* microsatellite loci amplified for paternity analyses. The annealing temperature is given as T_a_ and the number of alleles is given as N_a_

| Locus | Primer Sequence (5' - 3') | Repeat | T_a_ (°C) | Size | N_a_ |
| --- | --- | --- | --- | --- | --- |
| E7 | CCGGTTGCCCTTTGTCAAAG CTCGTCCCGCAAGAGGAAC | ACG | 60 | 147 - 174 | 5 |
| E14 | TCTCACCGCTATTACCATCTCT TGACTCGAGGTAGACGGAGA | ACT | 60 | 165 - 228 | 9 |
| E21 | AACGGAGATGGAGACGGAGA CGAAGCGCCACAGAATAACG | AAG | 60 | 124 - 145 | 6 |
| E22 | AGCCTGAAAACCCACCAACA GCAGAAGTCCGTTTCCTGGA | AAG | 60 | 152 - 179 | 7 |
| E24 | TTTCGTGAGTCAGAGACGCG CGGCAGAGAAGGAGGAATCG | AAG | 60 | 155 - 182 | 8 |
| E26 | TTTCTCCTCCGTCGTCGTTG CACGAGGAGCGAACGAGTAA | AAG | 60 | 133 - 157 | 8 |
| E27 | ACGACGGAACAACGGAGAAA TCAAGCCGAGCGAATGTAGT | ATC | 60 | 197 - 209 | 4 |
| E31 | AGTTTGATGCACGCGAAACC GCTCGCCCTCTTTCTCTCTT | ACT | 60 | 162 - 192 | 9 |
| E32 | GCCGTTCCAGCCAAGAGATA ACGTGCCGCAGTTTCGTATA | ACG | 60 | 138 - 153 | 6 |
| E33 | AGGACTGCCGGTTTAATGGT CAGCCACGCGTTCGATAAAT | AAGG | 60 | 199 - 235 | 8 |

Image S1

Experimental cage made out of shade cloth to provide forest understory conditions at the Fort Lauderdale Research and Education Center, Davie, FL.

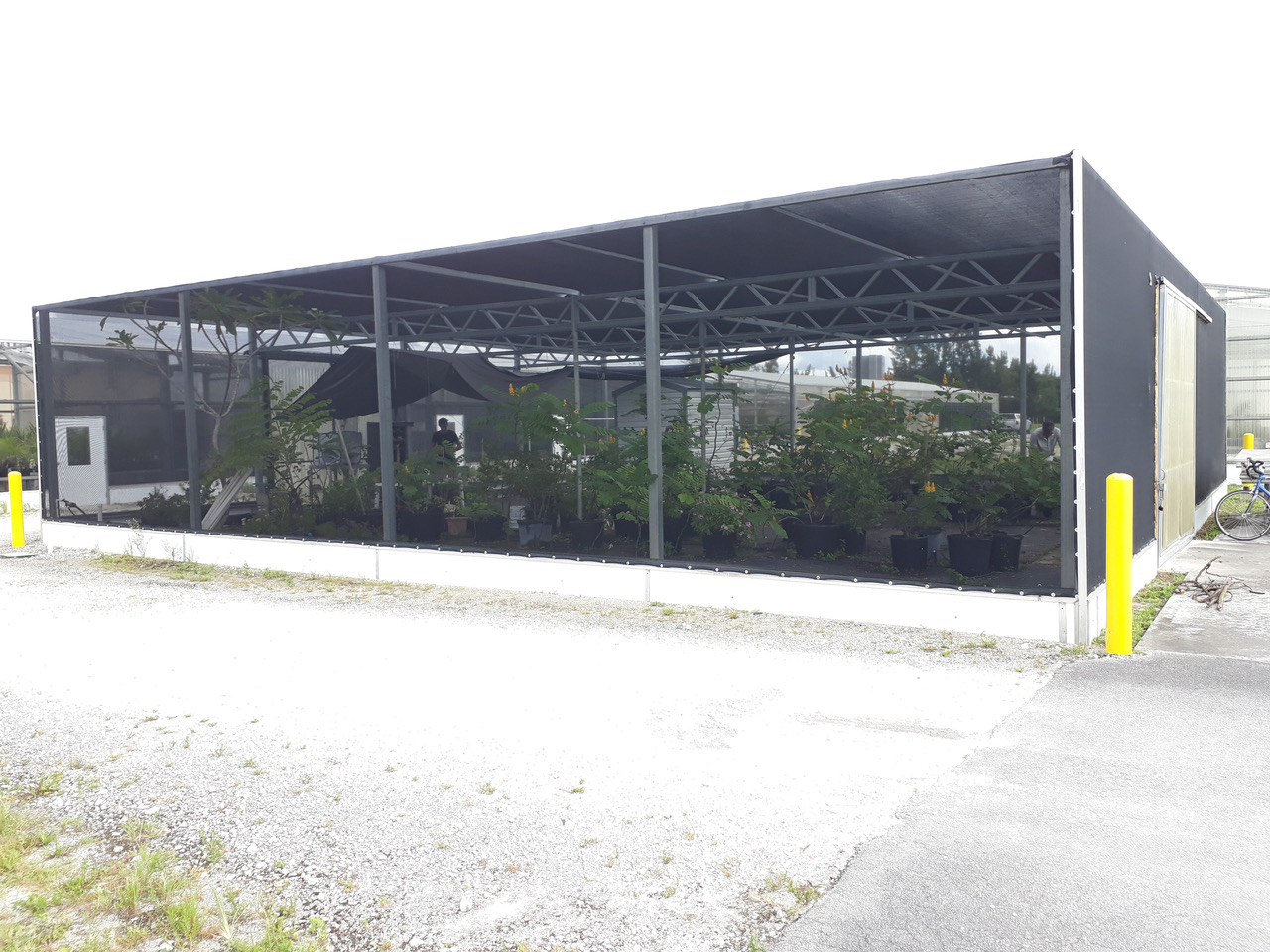

Image S2

Entrances to nesting boxes used by female Euglossa dilemma nesting in the cage. Wooden nest boxes were placed in a small shed inside the cage. Female bees accessed boxes through plastic tubes.

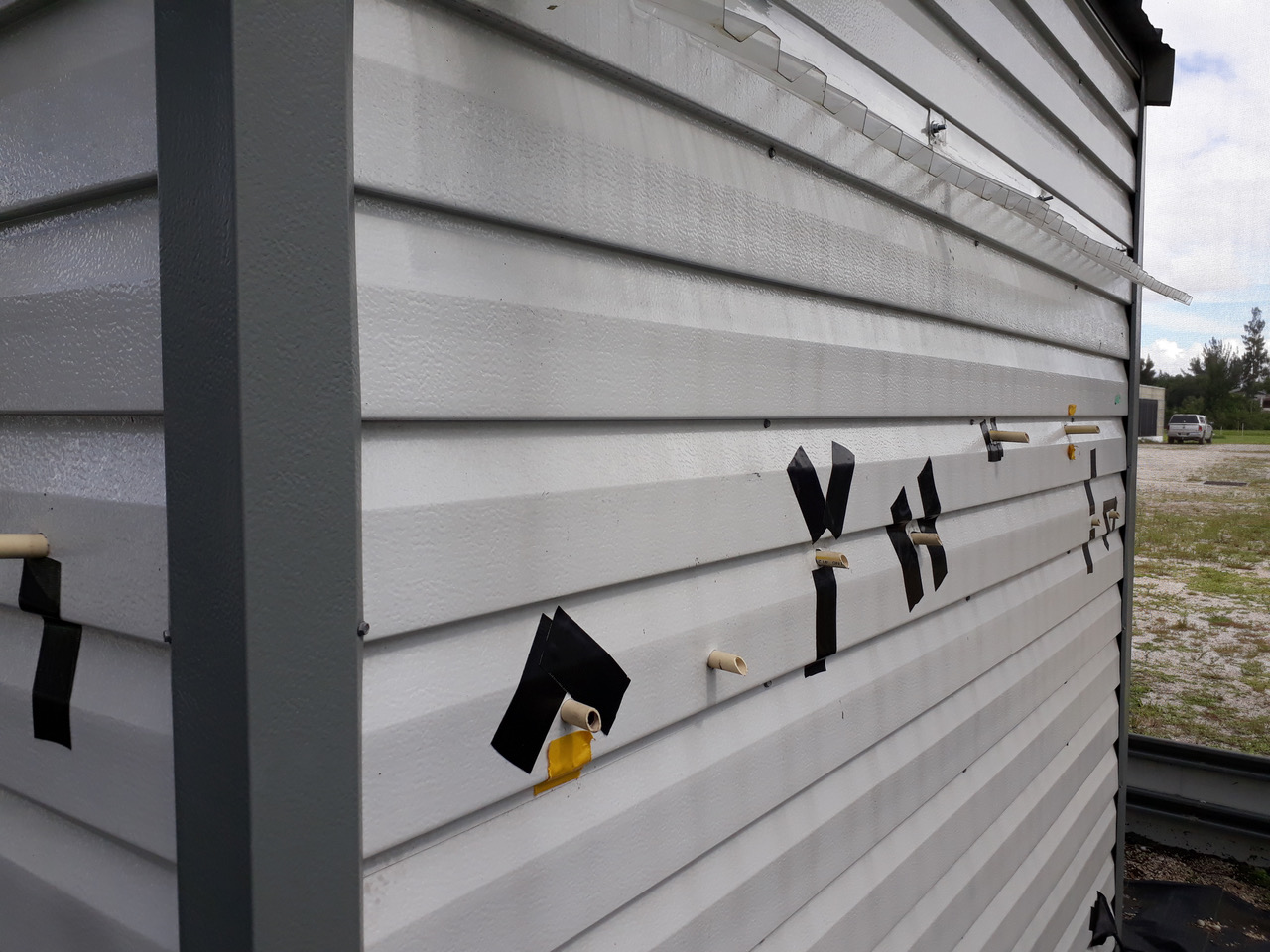

Image S3

Inside view of the cage with flowering food plants.
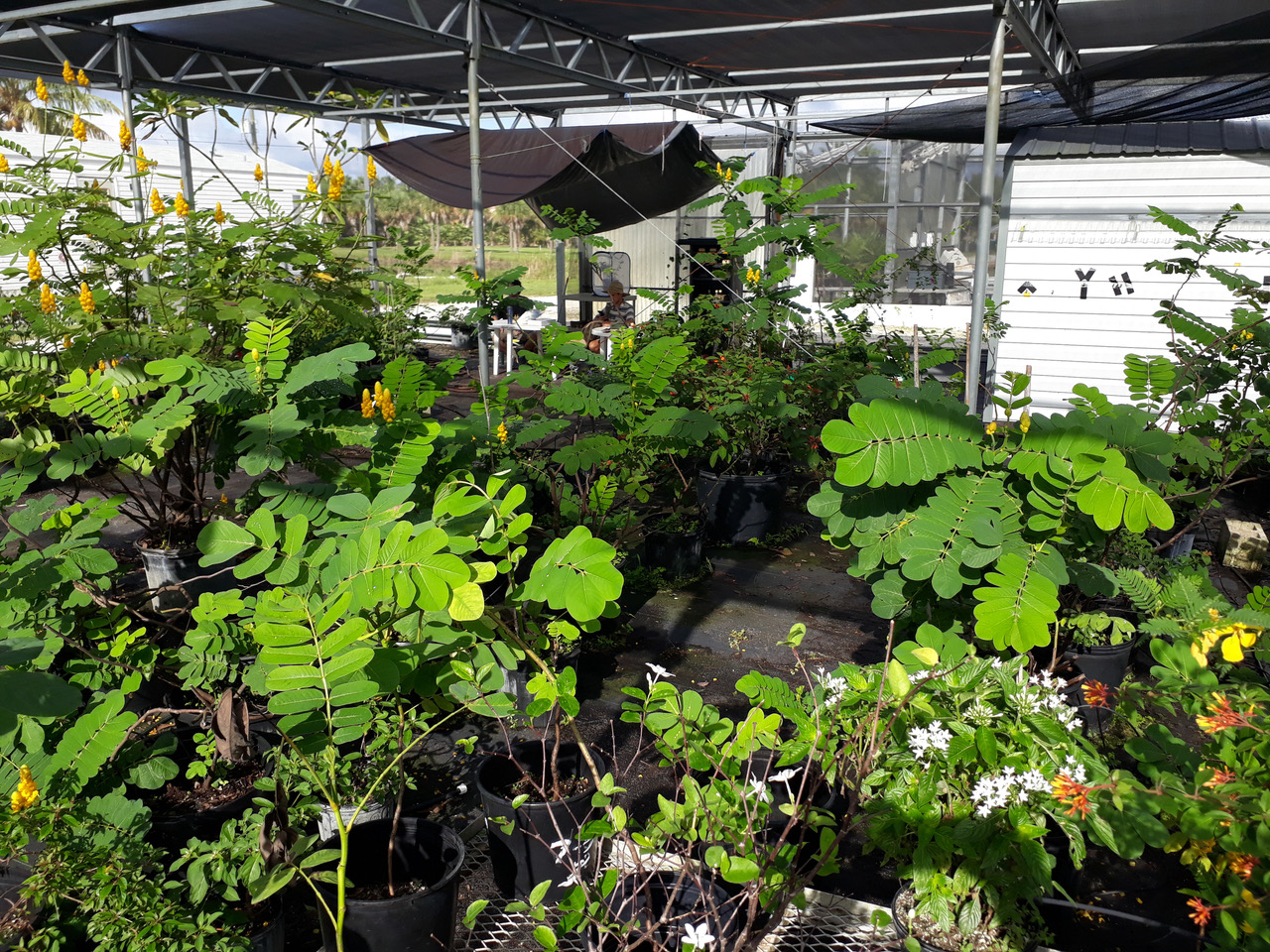

Movie S1.

**Orchid bee (*Euglossa dilemma*) copulation during experiment.** Note the high number of leg crossings performed by the male directly before mating takes place. Leg crossing behavior leads to perfume release during display. When male descents directly before copulation the everted genitals can be seen. Video taken by Jonas Henske.

Movie S2.

**Perfume harvest from wild caught bee using microcapillary.** Video taken by Thomas Eltz.

Movie S3.

**Perfume application of treatment male before introduction to the cage.** The perfume is deposited by a 1.2µL-CTC-syringe (Hamilton) to the hind-leg pocket. Note how the liquid is passively drawn inside after application. Video taken by Jonas Henske.
